## Supplementary Data for "Structure of the Mammalian Mediator"

### METHODS

**Purification of mMED for EM studies.** Mouse Mediator was immunopurified from nuclear extracts prepared from CH12 B lymphoma cell lines in which MED19 or MED25 were 3xFLAG-tagged at the N-terminus using CRISPR/Cas9[14]. Additional CRISPR/Cas9 editing was employed to create Mediator subunit truncations in the MED25 FLAG-tagged cell line. Med14 truncation was achieved by dual sgRNA transfection and screening clones for the desired in frame deletion. C-terminal Maltose-Binding-Protein (MBP) tagging of Med15 was performed by overexpression Med15-MBP in Med19 FLAG Med15 knock-out cells[14]. Sequences of sgRNAs and primers for cloning of targeting constructs are listed in Extended Data Table 3. Cell transfection and MBP overexpression was performed as previously described[14]. CH12 cell cultures were expanded to several billion cells in spinner flasks and nuclear extracts were prepared using a published protocol[14]. For mMED purification, nuclear extract was incubated with FLAG M2 agarose resin that had been pre-equilibrated in a buffer containing 50 mM HEPES pH 7.9, 300 mM KOAC, 1 mM EDTA, 10% Glycerol, and 0.2% NP-40. After incubation the resin was extensively washed with a buffer containing 50 mM HEPES pH 7.9, 300 mM KOAc, 1 mM EDTA, 10% Glycerol, 0.2% NP-40 and 1X mammalian protease inhibitor cocktail (Sigma P8340). This was followed by a second round of washing with a buffer containing 50 mM HEPES pH 7.9, 100 mM KCl, 1 mM EDTA, 5% Glycerol and 0.05% NP-40 and no protease inhibitors. Bound mMED was eluted with a buffer containing 500 µg/ml FLAG peptide (Sigma) in 50 mM HEPES pH 7.9, 100 mM KCl, 5% glycerol, 1 mM EDTA and 0.005% NP-40. Purified mMED fractions were flash-frozen in liquid nitrogen and stored at -80°C until needed for EM studies.

**mMED cryo-EM sample preparation, imaging and analysis.** MED19- or MED25-FLAG mMED cryo-EM samples were prepared on lacey carbon grids covered with a thin layer of continuous amorphous carbon (Ted Pella 01824). To prepare cryo-EM samples, purified mMED aliquots were concentrated 20-40 fold using a Vivaspin 500 centrifugal concentrator (Sartorius) and 2.5µL of purified mMED (100-250µg/mL) were pipetted onto grids plasma-cleaned for 6 sec on a Solarus plasma cleaner (Gatan) using an Ar/O2 gas mixture. Vitrification was performed in liquid ethane using a manual plunge-freeze apparatus. Imaging was performed on Talos Arctica transmission electron microscope (a Thermo Fisher) outfitted with an X-FEG electron source and operating at an acceleration voltage of 200kV. Automated data collection was carried out using stage-shift targeting in Leginon[30] and four separate image datasets were recorded using a K3 Summit direct electron detector (Gatan).

Information about imaging conditions and EM data collection statistics for mMED cryo-EM specimens is summarized in Extended Data Table 1. Cryo-EM movies from both zero-tilt and tilted (20°-40°) cryo-EM specimens were recorded to counteract the effect of anisotropic distribution of mMED particle orientations. Image processing was carried out using the Cryosparc[31] image processing package. Briefly, detector movie frames were subject to patch alignment and patch CTF refinement, followed by automated template-based particle picking. Repeated rounds of 2D image clustering were used to clean the initial image datasets. After this initial cleaning an *ab-initio* volume was calculated from each dataset and alignment parameters for cryo-EM images were obtained by 3D refinement. Images in each dataset were further screened by 3D image classification and the best images from each dataset were combined. Further rounds of 2D clustering and 3D classification of the combined dataset resulted in a selected set of images that were used for calculation of the final mMED cryo-EM map (Extended Data Figs 1-3).

**mMED cryo-EM map interpretation.** Map visualization and interpretation were done using Coot[32] for atomic model building, Phenix[33] for atomic model refinement and Chimera[34] for map visualization. Structure predictions for all mMED subunits were done using Phyre2[35] and I-TASSER[36]. Docking of the *S pombe* Mediator X-ray models derived from cryo-EM and X-ray studies (PDB accession codes 5U0P, 5U0S, and 5N9J), and information about Mediator subunit localization in yeast[9,10] and mammalian[14] Mediators guided initial watershed segmentation (with Chimera) of the mMED cryo-EM map. Atomic model building for mMED subunits was aided by consideration of available yeast Mediator subunit structures and structure prediction results. Atomic model building for Tail model subunits was done *de novo* for all subunits, except for MED23, for which the X-ray structure of human MED23 (PDB 6H02) was used as a starting point. Phenix was used for refinement of individual subunit atomic models and for refinement of the overall mMED atomic model (Extended Data Table 1).

**Localization of mMED subunit domains by EM analysis of truncation and MBP-labeled mutants.** Localization of mMED subunit domains through truncation or MBP-tagging was done by EM image analysis of mMED particles preserved in stain. Stained samples were prepared on continuous carbon EM grids (EMS 017543) and preserved using 2% uranyl acetate. Stained samples were imaged using a Talos L120C transmission electron microscope (Thermo-Fisher) outfitted with a $LaB_6$ filament and operating at an acceleration voltage of 120kV. Automated data collection was carried out using Leginon[30], with images recorded using a Ceta CMOS detector at a magnification of 36,000X (corresponding to a pixel size of 3.98Å). Particles were automatically picked from micrographs with DogPicker[37], using a low threshold to capture all mMED particle images. Roughly 20,000 particle images picked for each experiment were subject to image clustering using ISAC[38]. The cleanest mMED averages were selected and their intensities were normalized to an average density of zero and a standard deviation of 1.0. Multiple average pairs were compared to identify average pairs that would result in the cleanest difference maps. Difference and heat maps were calculated and displayed using a custom image processing and plotting script written in Matlab, which allowed for interactive fine-tuning of cross-correlation-based map alignment prior to difference map calculation. Difference maps were also visualized as heat maps colored by standard deviation, to facilitate their interpretation.

**Data availability**

Cryo-EM maps and atomic coordinates have been deposited with the Electron Microscopy Data Bank (with accession code EMD-21514 and Protein Data Dank (accession code PDB ID 6W1S). The accession numbers for the deep-sequencing data reported in this paper are found at GSE145902.

### EXTENDED DATA TABLES

#### Extended Data Table 1. Cryo-EM data collection, processing and model validation statistics

| **Data collection and Processing** | |
|---|---|
| Microscope | Talos Arctica (ThermoFisher) |
| Detector | K3 Summit (Gatan) |
| Voltage (keV) | 200 |
| Defocus range (□m) | 0.8 - 3.5 |
| Number of movies (0°/20°/30°/40° tilt) | 2,705/6,464/2,928/1,495 (13,592 total) |
| Frames per movie | 40 - 60 |
| Exposure time per frames (ms) | 75 - 100 |
| Magnification | 36,000x |
| Pixel size (Å) | 1.11 |
| Dose rate ($e^-$/pixel/sec) | 19 - 22 |
| Total dose per movie ($e^-$/Å$^2$) | 50 - 80 |
| Initial number of particle images | 2,319,481 |
| Final number of particle images | 217,557 |
| Particle symmetry | C1 |
| Overall map resolution (Å) | 4.0 |
| FSC threshold | 0.143 |
| Map resolution range (Å) | 3.4 - >10 |
| Directional resolution range (Å) | 3.4 – 6.8 |
| Sphericity of 3D FSC | 0.838 |
| Map sharpening B-factor (Å$^2$) | 74 |
| **Atomic model validation statistics** | |
| **Model Composition** | |
| Protein Chains | 25 |
| Protein Residues | 7,434 |
| Non-hydrogen atoms | 47,305 |
| **R.M.S deviations** | |
| Bond lengths (Å) | 0.003 |
| Bond angles (°) | 0.682 |
| **Validation** | |
| MolProbity score | 2.07 |
| Clashscore | 8.11 |
| Poor rotamers (%) | 0 |
| **Ramachandran plot** | |
| Favored (%) | 86.86 |
| Allowed (%) | 13.02 |
| Disallowed (%) | 0.13 |

**Extended Data Table 2: Subunit module assignment and atomic model information**

| MmMED Subunit | Chain ID | Module | Color | # of Residues | Residues in atomic model (%) |
|---|---|---|---|---|---|
| MED1 | A | Middle | purple | 1575 | ~1-520 (33) |
| MED4 | B | Middle | light blue | 270 | 30-129 (37) |
| MED6 | C | Head | yellow | 246 | 9-246 (97) |
| MED7 | D | Middle | sandy brown | 233 | 11-167 (67) |
| MED8 | E | Head | forest green | 268 | 7-186 (67) |
| MED9 | F | Middle | hot pink | 142 | 61-136 (53) |
| MED10 | G | Middle | magenta | 135 | 61-130 (52) |
| MED11 | H | Head | orange red | 117 | 8-115 (92) |
| MED14 | I | - | green | 1459 | 149-975/1172-1450 (76) |
| MED15 | J | Tail | red | 789 | 620-787 (21) |
| MED16 | K | Tail | khaki | 828 | 3-825 (99) |
| MED17 | L | Head | dodger blue | 649 | 15-645 (97) |
| MED18 | M | Head | tan | 208 | 17-203 (90) |
| MED19 | N | Middle | navy blue | 244 | 78-136 (24) |
| MED20 | O | Head | cyan | 212 | 3-200 (93) |
| MED21 | P | Middle | dark green | 144 | 3-128 (88) |
| MED22 | Q | Head | medium purple | 200 | 9-139 (66) |
| MED23 | R | Tail | plum | 1367 | 1-1334 (98) |
| MED24 | S | Tail | dark cyan | 987 | 4-985 (99) |
| MED25 | T | Tail | salmon | 745 | 5-216 (28) |
| MED26 | U | Middle-Head | | 588 | Not included |
| MED27 | V | Tail | violet red | 311 | 8-304 (95) |
| MED28 | W | Tail | goldenrod | 178 | 32-149 (66) |
| MED29 | X | Tail | blue | 199 | 52-185 (67) |
| MED30 | Y | Tail | olive drab | 178 | 27-178 (85) |
| MED31 | Z | Middle | firebrick | 131 | 13-105 (71) |

**Extended Data Table 3: Primer and cloning strategy for cell line creation.** Excel Document.

### EXTENDED DATA MOVIE

**Extended Data Movie 1:** A movie showing the mMED atomic model, including a close up view the Tail module.

a

MED19-FLAG

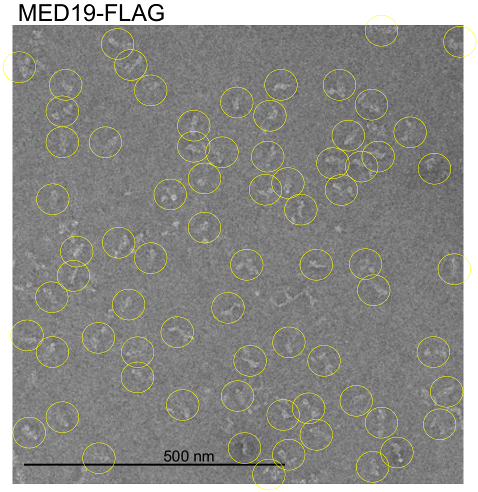

MED25-FLAG

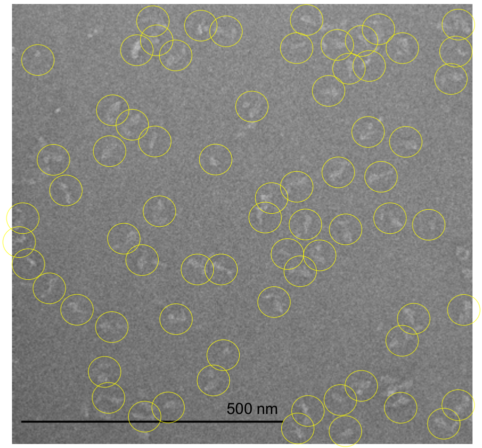

b

| Identified Proteins | Accession Identifier | Molecular Weight | MED19-FLAG Peptide Counts |
|---|---|---|---|
| Mediator of RNA polymerase II transcription subunit 12 | MED12_MOUSE | 245 kDa | 290 |
| Mediator of RNA polymerase II transcription subunit 14 | MED14_MOUSE | 161 kDa | 276 |
| Actin, cytoplasmic 2 | ACTG_MOUSE | 42 kDa | 241 |
| Mediator of RNA polymerase II transcription subunit 13-like | MD13L_MOUSE | 242 kDa | 224 |
| Mediator of RNA polymerase II transcription subunit 23 | MED23_MOUSE | 156 kDa | 220 |
| DNA-directed RNA polymerase II subunit RPB1 | RPB1_MOUSE | 217 kDa | 216 |
| Mediator of RNA polymerase II transcription subunit 1 | MED1_MOUSE | 167 kDa | 163 |
| Mediator of RNA polymerase II transcription subunit 24 | MED24_MOUSE | 110 kDa | 159 |
| Mediator of RNA polymerase II transcription subunit 16 | MED16_MOUSE | 92 kDa | 153 |
| DNA-directed RNA polymerase II subunit RPB2 | RPB2_MOUSE | 134 kDa | 136 |
| Mediator of RNA polymerase II transcription subunit 17 | MED17_MOUSE | 72 kDa | 132 |
| Myosin-9 | MYH9_MOUSE | 226 kDa | 124 |
| Serine/threonine-protein kinase 38 | STK38_MOUSE | 54 kDa | 117 |
| Unconventional myosin-Ie | MYO1E_MOUSE | 127 kDa | 99 |
| Actin, alpha cardiac muscle 1 | ACTC_MOUSE | 42 kDa | 91 |
| Mediator of RNA polymerase II transcription subunit 15 | MED15_MOUSE | 87 kDa | 87 |
| Gelsolin | GELS_MOUSE | 86 kDa | 82 |
| Mediator of RNA polymerase II transcription subunit 27 | MED27_MOUSE | 35 kDa | 78 |
| Mediator of RNA polymerase II transcription subunit 4 | MED4_MOUSE | 30 kDa | 67 |
| Tropomodulin-3 | TMOD3_MOUSE | 40 kDa | 67 |
| Unconventional myosin-XVIIIa | MY18A_MOUSE | 233 kDa | 66 |
| Advillin | AVIL_MOUSE | 92 kDa | 62 |
| Beta-actin-like protein 2 | ACTBL_MOUSE | 42 kDa | 62 |
| Mediator of RNA polymerase II transcription subunit 8 | MED8_MOUSE | 29 kDa | 57 |
| Mediator of RNA polymerase II transcription subunit 7 | MED7_MOUSE | 27 kDa | 53 |
| Mediator of RNA polymerase II transcription subunit 26 | MED26_MOUSE | 65 kDa | 52 |
| Mediator of RNA polymerase II transcription subunit 19 | MED19_MOUSE | 26 kDa | 52 |
| Mediator of RNA polymerase II transcription subunit 10 | MED10_MOUSE | 16 kDa | 48 |
| DNA-directed RNA polymerase II subunit RPB3 | RPB3_MOUSE | 31 kDa | 46 |
| Mediator of RNA polymerase II transcription subunit 20 | MED20_MOUSE | 23 kDa | 46 |
| Mediator of RNA polymerase II transcription subunit 6 | MED6_MOUSE | 28 kDa | 45 |
| Cyclin-dependent kinase 8 | CDK8_MOUSE | 53 kDa | 42 |
| Mediator of RNA polymerase II transcription subunit 11 | MED11_MOUSE | 13 kDa | 42 |
| Mediator of RNA polymerase II transcription subunit 21 | MED21_MOUSE | 16 kDa | 35 |
| DNA-directed RNA polymerase II subunit RPB7 | RPB7_MOUSE | 19 kDa | 35 |
| Mediator of RNA polymerase II transcription subunit 28 | MED28_MOUSE | 20 kDa | 34 |
| Mediator of RNA polymerase II transcription subunit 30 | MED30_MOUSE | 20 kDa | 34 |
| Mediator of RNA polymerase II transcription subunit 22 | MED22_MOUSE | 22 kDa | 33 |
| Cyclin-dependent kinase 19 | CDK19_MOUSE | 57 kDa | 33 |
| Thyroid hormone receptor-associated protein 3 | TR150_MOUSE | 108 kDa | 30 |
| Mediator of RNA polymerase II transcription subunit 18 | MED18_MOUSE | 24 kDa | 28 |
| Unconventional myosin-Ic | MYO1C_MOUSE | 122 kDa | 28 |
| DNA-directed RNA polymerases I, II, and III subunit RPABC1 | RPAB1_MOUSE | 25 kDa | 27 |
| Mediator of RNA polymerase II transcription subunit 29 | MED29_MOUSE | 21 kDa | 27 |
| F-actin-capping protein subunit beta | CAPZB_MOUSE | 31 kDa | 27 |
| Unconventional myosin-Ig | MYO1G_MOUSE | 117 kDa | 27 |
| Mediator of RNA polymerase II transcription subunit 31 | MED31_MOUSE | 16 kDa | 25 |
| DNA-directed RNA polymerase II subunit RPB9 | RPB9_MOUSE | 15 kDa | 25 |
| Mediator of RNA polymerase II transcription subunit 9 | MED9_MOUSE | 16 kDa | 24 |
| Actin-related protein 3 | ARP3_MOUSE | 47 kDa | 24 |
| Myosin regulatory light chain 12B | ML12B_MOUSE | 20 kDa | 23 |
| Probable helicase senataxin | SETX_MOUSE | 298 kDa | 22 |
| Myosin light polypeptide 6 | MYL6_MOUSE | 17 kDa | 20 |
| Calmodulin | CALM_MOUSE | 17 kDa | 19 |
| Cyclin-C | CCNC_MOUSE | 33 kDa | 18 |
| Heat shock cognate 71 kDa protein | HSP7C_MOUSE | 71 kDa | 17 |
| DNA-directed RNA polymerase II subunit RPB4 | RPB4_MOUSE | 16 kDa | 17 |
| Mediator of RNA polymerase II transcription subunit 25 | MED25_MOUSE | 78 kDa | 17 |
| F-actin-capping protein subunit alpha-1 | CAZA1_MOUSE | 33 kDa | 17 |
| Keratin, type I cytoskeletal 10 | K1C10_MOUSE | 58 kDa | 17 |
| Actin-related protein 2 | ARP2_MOUSE | 45 kDa | 15 |
| DNA-directed RNA polymerases I, II, and III subunit RPABC3 | RPAB3_MOUSE | 17 kDa | 14 |
| Unconventional myosin-Va | MYO5A_MOUSE | 216 kDa | 13 |
| Tubulin beta-5 chain | TBB5_MOUSE | 50 kDa | 13 |
| Unconventional myosin-Id | MYO1D_MOUSE | 116 kDa | 12 |
| DNA-directed RNA polymerase II subunit RPB11 | RPB11_MOUSE | 13 kDa | 11 |
| DNA-directed RNA polymerases I, II, and III subunit RPABC5 | RPAB5_MOUSE | 8 kDa | 11 |
| F-actin-capping protein subunit alpha-2 | CAZA2_MOUSE | 33 kDa | 11 |
| Actin-related protein 2/3 complex subunit 1B | ARC1B_MOUSE | 41 kDa | 11 |
| Actin-related protein 2/3 complex subunit 4 | ARPC4_MOUSE | 20 kDa | 11 |
| Enhancer of rudimentary homolog | ERH_MOUSE | 12 kDa | 10 |
| Keratin, type II cytoskeletal 1 | K2C1_MOUSE | 66 kDa | 10 |
| Actin-related protein 2/3 complex subunit 2 | ARPC2_MOUSE | 34 kDa | 10 |
| Plastin-2 | PLSL_MOUSE | 70 kDa | 10 |
| Keratin, type II cytoskeletal 5 | K2C5_MOUSE | 62 kDa | 9 |
| Keratin, type I cytoskeletal 13 | K1C13_MOUSE | 48 kDa | 9 |
| Keratin, type I cytoskeletal 14 | K1C14_MOUSE | 53 kDa | 8 |
| Keratin, type II cytoskeletal 73 | K2C73_MOUSE | 59 kDa | 8 |
| Bcl-2-associated transcription factor 1 | BCLF1_MOUSE | 106 kDa | 8 |
| Myosin light chain kinase 2, skeletal/cardiac muscle | MYLK2_MOUSE | 66 kDa | 7 |
| 78 kDa glucose-regulated protein | GRP78_MOUSE | 72 kDa | 6 |
| Actin-related protein 2/3 complex subunit 3 | ARPC3_MOUSE | 21 kDa | 6 |
| Spindlin-1 | SPIN1_MOUSE | 30 kDa | 5 |
| Gamma-adducin | ADDG_MOUSE | 79 kDa | 5 |
| Alpha-adducin | ADDA_MOUSE | 81 kDa | 4 |
| Tubulin alpha-1B chain | TBA1B_MOUSE (+1) | 50 kDa | 4 |
| Lymphocyte-specific protein 1 | LSP1_MOUSE | 37 kDa | 4 |
| E3 ubiquitin-protein ligase TRIM21 | RO52_MOUSE | 54 kDa | 4 |
| Elongation factor 1-alpha 1 | EF1A1_MOUSE | 50 kDa | 3 |
| Actin-related protein 2/3 complex subunit 5-like protein | ARP5L_MOUSE | 17 kDa | 3 |
| Collagen alpha-2(I) chain | CO1A2_MOUSE | 130 kDa | 3 |
| Q3TTY5\|K22E_MOUSE-DECOY | Q3TTY5\|K22E_MOUSE- | ? | 2 |
| U4/U6 small nuclear ribonucleoprotein Prp4 | PRP4_MOUSE | 58 kDa | 2 |

Mediator Subunit
RNAPII Subunit
Kinase Module Subunit
Uncommon Contaminant

**Extended Data Figure 1. Stained images and MudPIT analysis of mMED preparations. a,** Images of MED19-FLAG and MED25-FLAG mMED preserved in stain show homogenous particles and absence of any non-Mediator contaminants. **b,** MudPIT analysis of purified mMED fractions shows that all Mediator subunits are present and that the only other proteins detected at any significant level were components of the kinase module and RNAPII subunits.

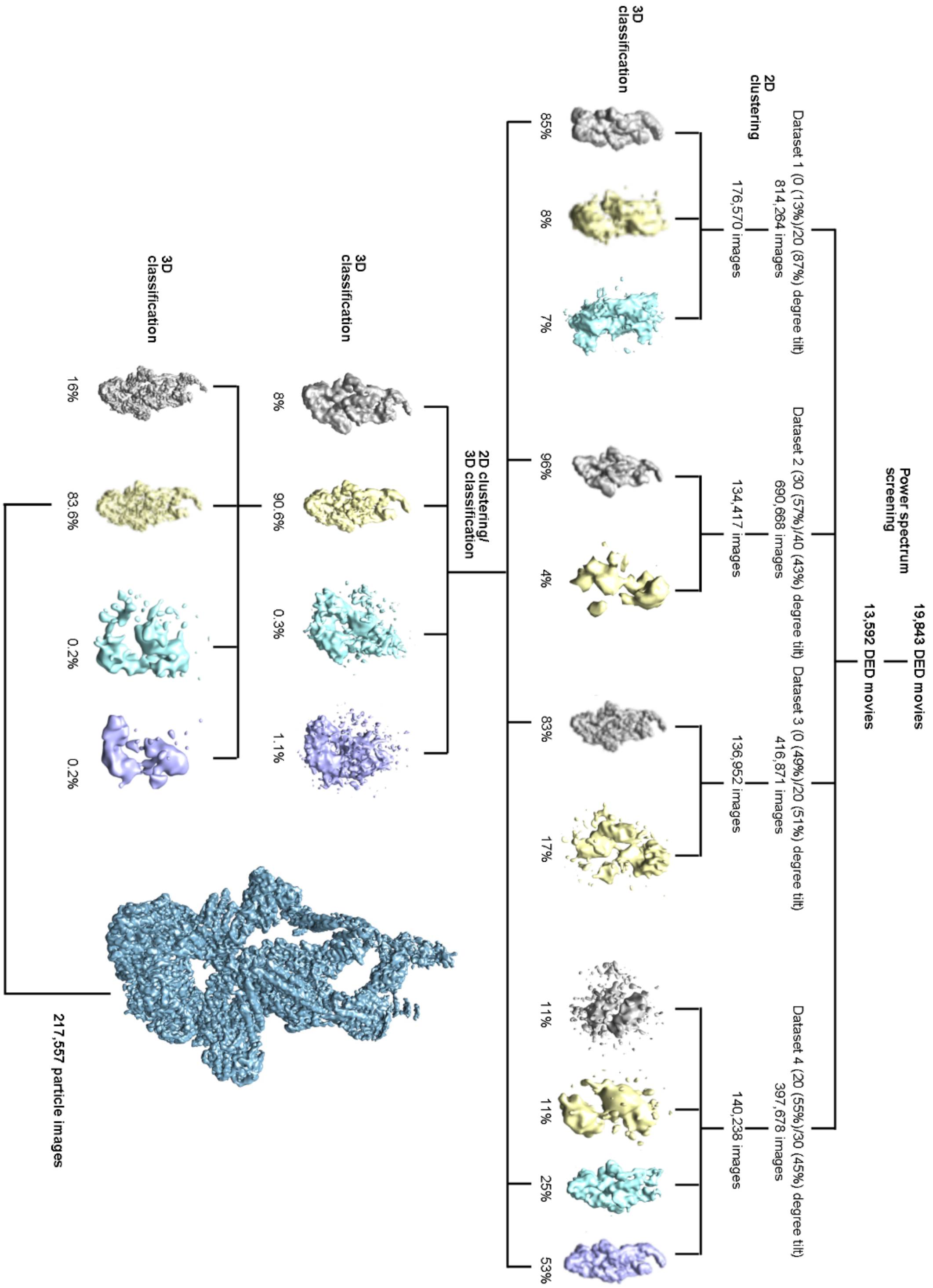

**Extended Data Figure 2. 2D clustering and 3D classification diagram for mMED cryo-EM analysis.** Sequence of power spectra screening, 2D clustering and 3D classification steps during selection and analysis of mMED cryo-EM particle images.

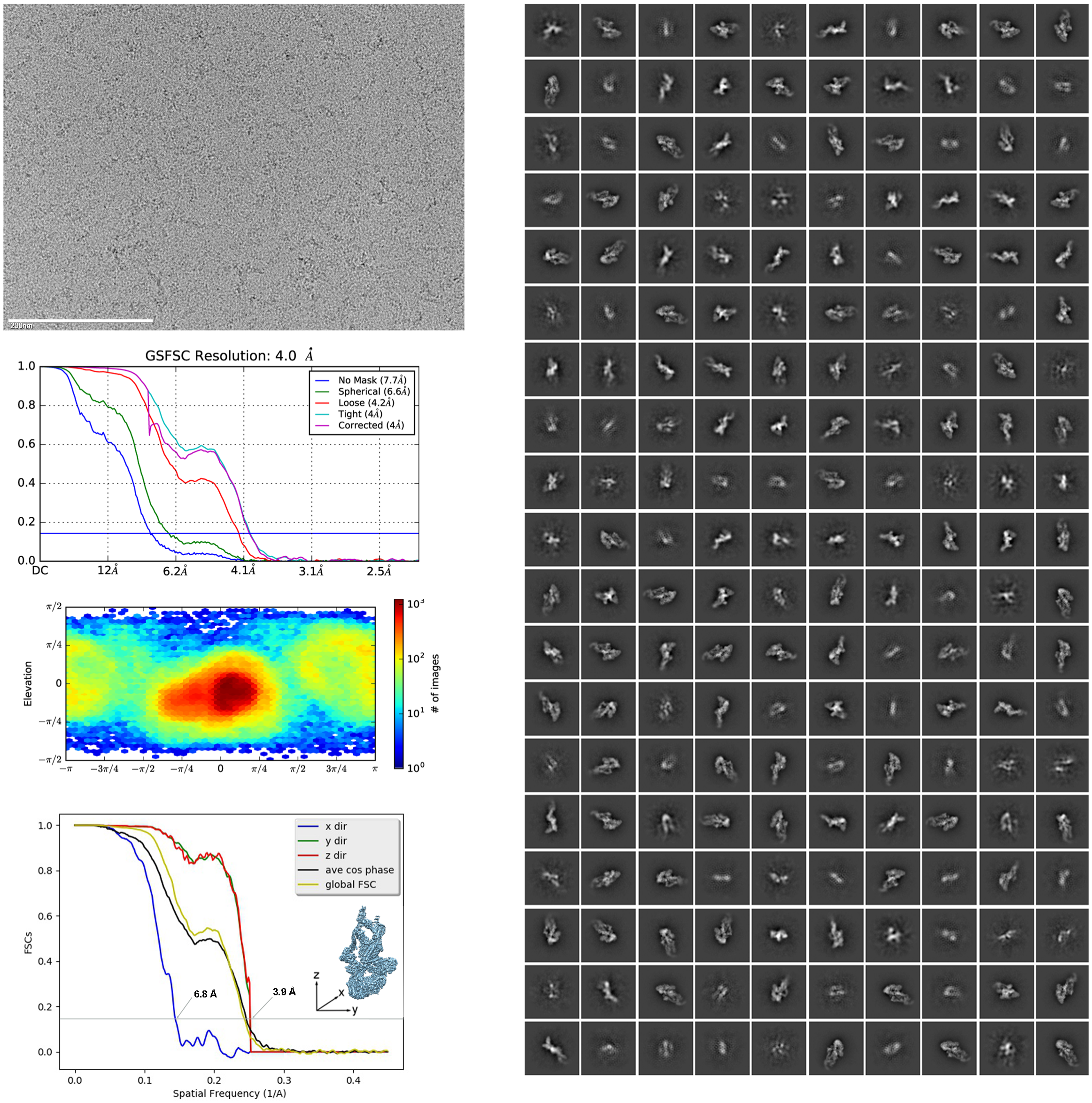

**Extended Data Figure 3. mMED cryo-EM analysis. a,** Typical mMED micrograph showing particles cryo-preserved on a thin carbon film (0° tilt). Scale bar 200nm. **b,** 2D class averages obtained from clustering of a combination of untilted and tilted mMED images. **c,** Fourier Shell Correlation (FSC) plot used to estimate the resolution of the mMED cryo-EM map to 4.0Å. **d,** Angular distribution plot for the final combined mMED cryo-EM dataset. **e,** Directional FSC plots showing the resolution of the mMED cryo-EM map along 3 perpendicular directions.

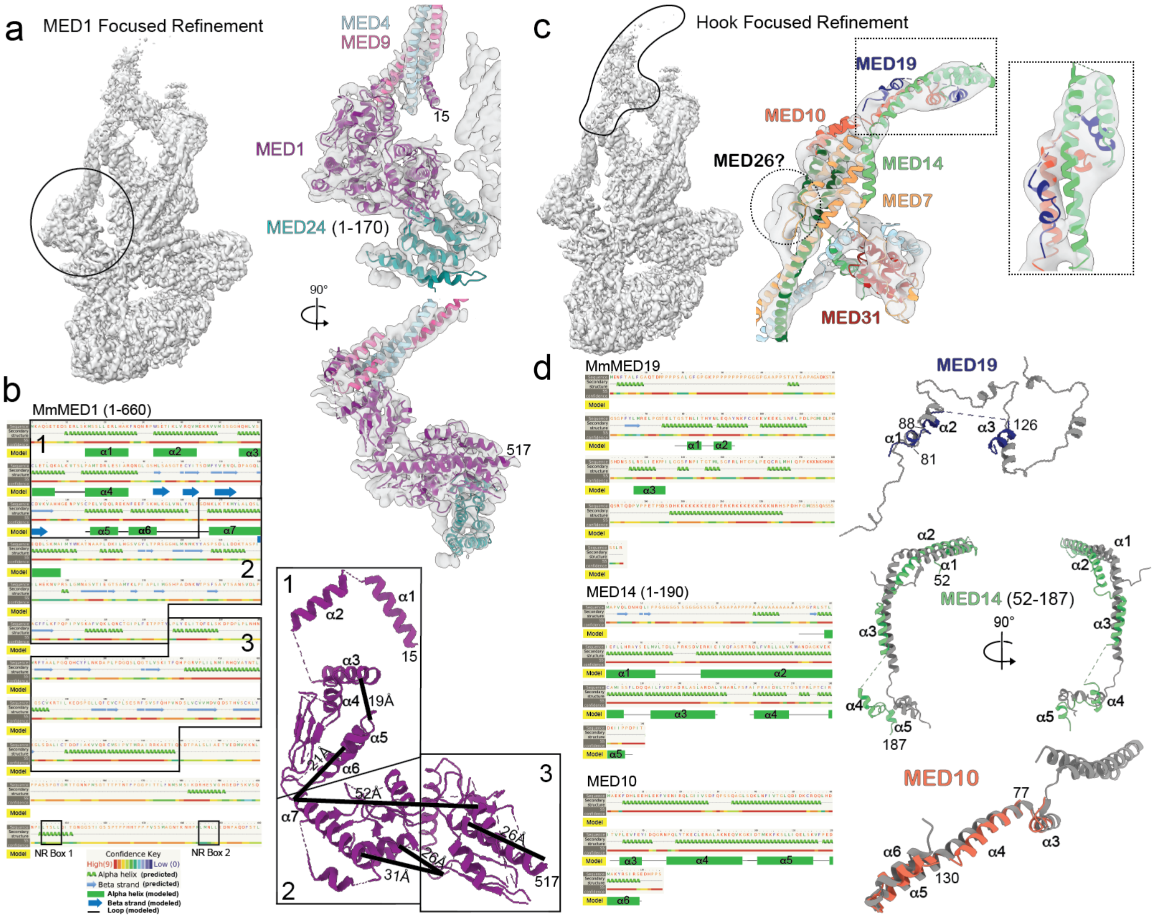
a
MED1 Focused Refinement
MED4
MED9
15
MED1
MED24 (1-170)
90°
517
b
MmMED1 (1-660)
1
2
3
α1
α2
α3
α4
α5
α6
α7
NR Box 1
NR Box 2
Confidence Key
High(9)
Low (0)
Alpha helix (predicted)
Beta strand (predicted)
Alpha helix (modeled)
Beta strand (modeled)
Loop (modeled)
19Å
52Å
26Å
31Å
c
Hook Focused Refinement
MED19
MED10
MED26?
MED14
MED7
MED31
d
MmMED19
MED14 (1-190)
MED10
MED19
88
81
126
MED14 (52-187)
52
187
90°
MED10
77
130

**Extended Data Figure 4. Focused refinement maps, secondary structure predictions and structure of the MED1 and hook portions of the Middle module. a,** Focused refinement map (left) including MED1 and neighboring portions of MED4, MED9, MED14 and MED24 and partial model of MED1 (~aa 1-520) showing its interaction with MED4-MED9 and MED24 (right). The MED1 N-terminus wraps around the bottom portion of MED4-MED9 (in light blue and pink, respectively) and a N-terminal α-helix contacts both subunits. From there, alternating $\beta$-strand and α-helical domains extend to contact the very N-terminal portion of MED24 (in dark cyan) through an extended $\beta$-sheet domain. With its alternating arrangement of helices and strands, the structure of the ordered N-terminal portion of MED1 is reminiscent of MED14 and could undergo conformational rearrangements allowing the Middle to move without breaking its contact with MED24. **b,** Secondary structure prediction for MED1 (left) calculated using Phyre2[35]. No ordered domains are predicted after the first ~720 aa. Three regions, delineated as indicated by black outlines, correspond to portions of the partial MED1 model (right). Region 3 includes $\beta$-strands that interact directly with a hydrophobic patch on the MED24 N-terminus. **c,** Focused refinement map (left) of the hook/knob portion of the Middle module and models of the corresponding subunits fitted into the map (right). **d,** Secondary structure predictions (left, from Phyre2[35]) for MED10, MED19 and MED14's 150 N-terminal residues show them to be mostly helical and to correspond both in size and expected secondary structure with corresponding *S pombe* Mediator subunits (MED19 is larger in mouse than yeast). Molecular models of mammalian hook subunits (MED14 N-terminus, MED10, MED19) based on secondary structure predictions and the focused refinement map show structures that match well corresponding portions of the atomic model of *S pombe* Mediator (PDB 5N9J).

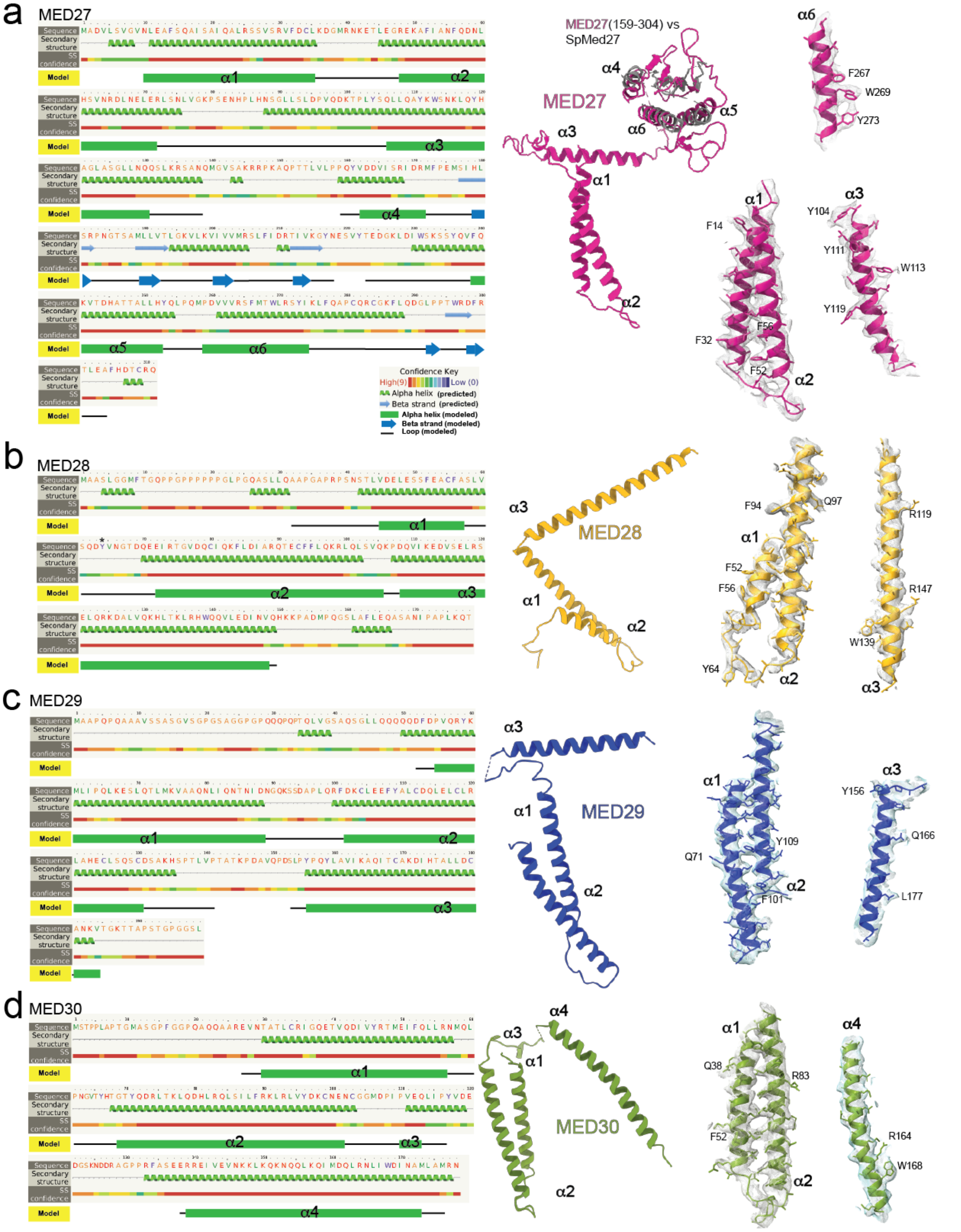
a
MED27
MED27(159-304) vs
SpMed27
MED27
α1
α2
α3
α4
α5
α6
F267
W269
Y273
F14
Y104
Y111
W113
Y119
F56
F32
F52
Confidence Key
High(9)
Low (0)
Alpha helix (predicted)
Beta strand (predicted)
Alpha helix (modeled)
Beta strand (modeled)
Loop (modeled)
b
MED28
MED28
F94
Q97
R119
F52
F56
R147
W139
Y64
c
MED29
MED29
Y156
Q166
Q71
Y109
F101
L177
d
MED30
MED30
Q38
R83
F52
R164
W168
Sequence
Secondary structure
SS confidence
Model

**Extended Data Figure 5. Interpretation and structure of upper Tail subunits. a-d,** Secondary structure predictions for MED27-30 (left) calculated using Phyre2[35] show that all 4 subunits (which are all very similar in size) are expected to have the same overall secondary structure, but that the length of specific $\alpha$-helices and loops differs from one subunit to another. This, and clear bulky side-chain densities in the corresponding portions of the mMED cryo-EM map (right) made possible building an accurate model of the upper Tail.

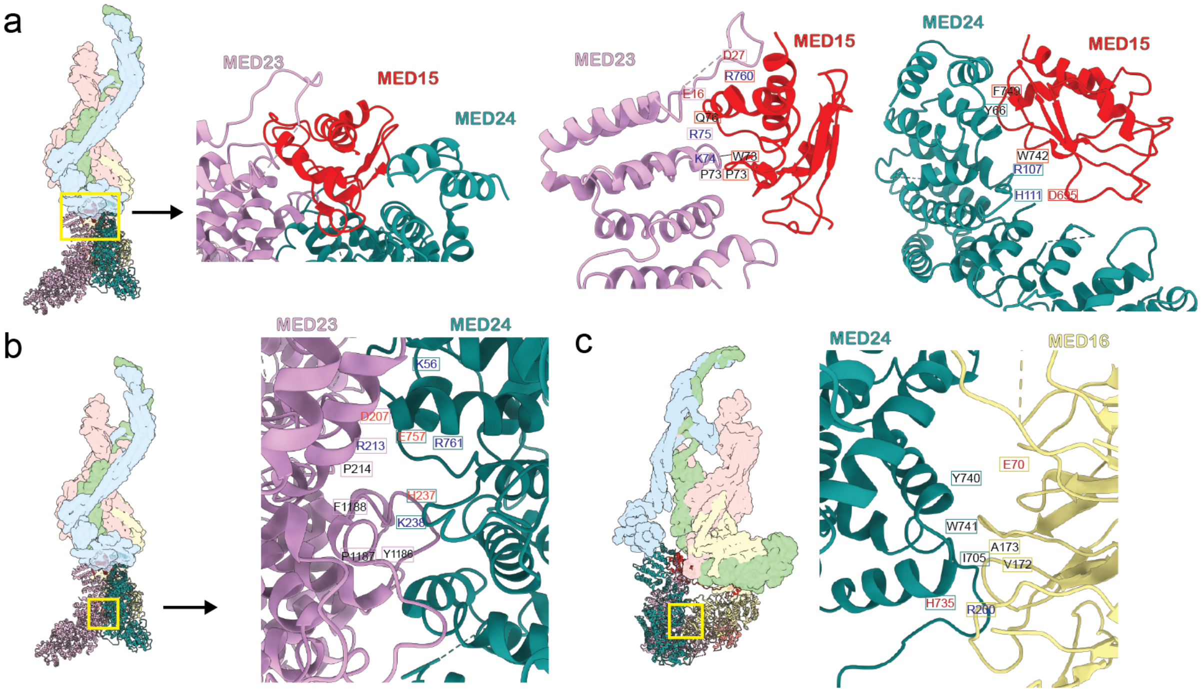

**Extended Data Figure 6. Inter-subunit contacts in the lower Tail.** Lower Tail subunit interfaces show a number of charged, hydrophobic and cationic-$\pi$ interactions, which are depicted by coloring the involved residues in black (polar and hydrophobic amino acids), red (negatively charged) and blue (positively charged). **a,** Interaction of MED15 with MED23 and MED24. The MED15/MED23 interface shows electrostatic interactions between MED23 D27, E16, R75 and MED15 R760 and Q763, with potential hydrogen-bonding between MED23 E16 and MED15 Q763. There is a cationic-$\pi$ interaction between MED23 K74 and MED15 W737, followed by hydrophobic-interaction between MED23 P73 and MED15 P735. The MED15/MED24 interface includes hydrophobic (MED24 Y66-MED15 F749), cationic-$\pi$ (MED24 R107-MED15 W742), and electrostatic (MED24 H111-MED15 D695) interactions. **b,** In the MED23/MED24 interface. MED23 D207 interacts with K56 and R761 on MED24 to form an electrostatic interface, with a potential salt-bridge between MED23 R213 and MED24 E757. Below this interaction, there is an aromatic cluster formed by MED23 P214, F1188, P1187, Y1186 and MED24 H237, with a cationic-$\pi$ interaction between MED23 F1188 and MED24 K238. **c,** At the MED16/MED24 interface, MED24 Y740 forms a hydrogen-bond with E70 of MED16. A hydrophobic patch between MED24 W741, I705 and MED16 A173, V172 is observed, as well as a charge interaction between MED24 H735 and MED16 R200.

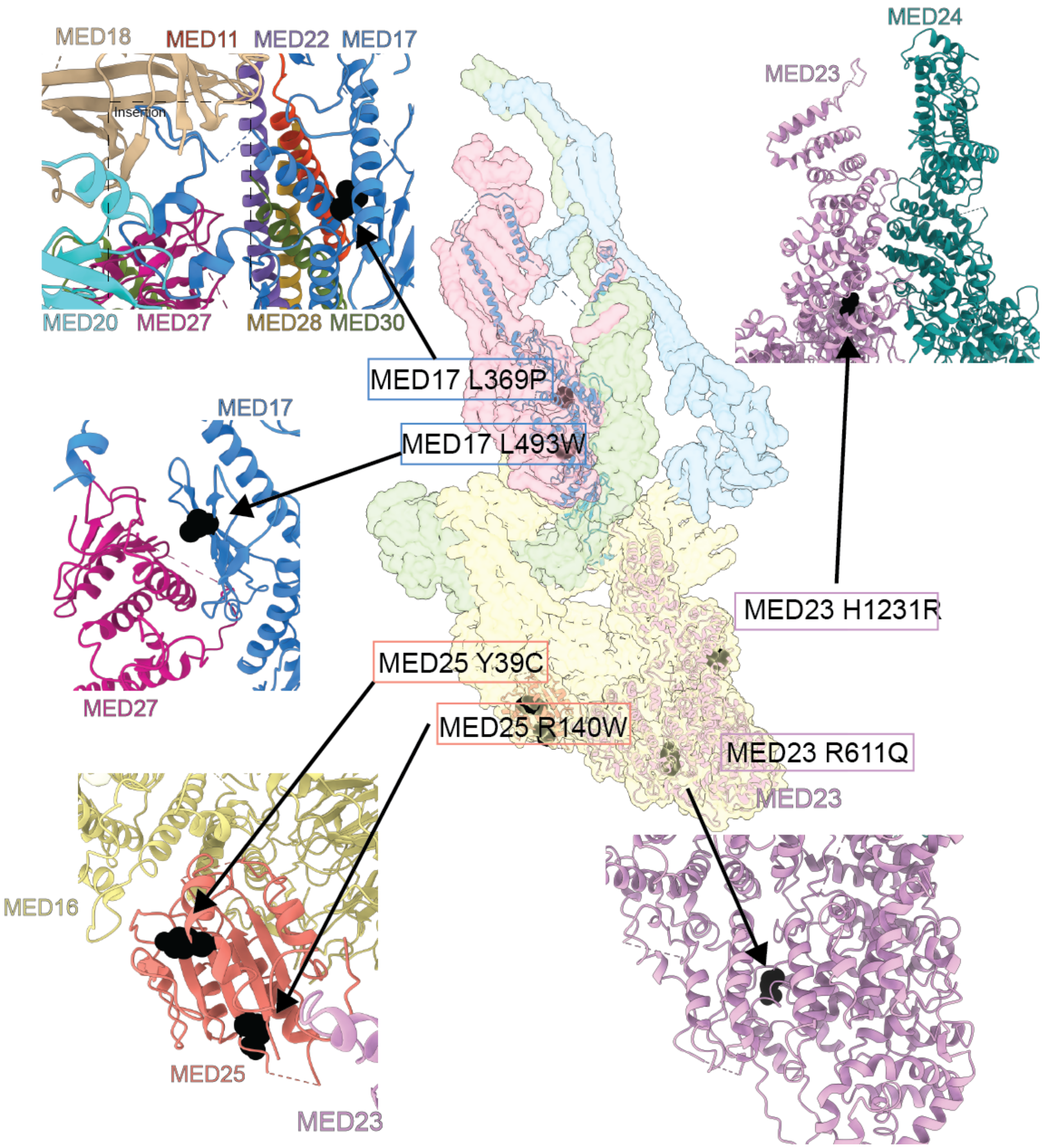

**Extended Data Figure 7. Disease-associated mMED mutations and mMED subunit IDRs. a,** Location of known disease-associated mammalian Mediator mutations.
